# Epigenetic Remodeling by Vitamin C Potentiates the Differentiation of Mouse and Human Plasma Cells

**DOI:** 10.1101/2021.09.15.460473

**Authors:** Heng-Yi Chen, Ana Alamonte-Loya, Fang-Yun Lay, Eric Johnson, Edahi Gonzalez-Avalos, Jieyun Yin, Qin Ma, Daniel J. Wozniak, Fiona E. Harrison, Chan-Wang Jerry Lio

## Abstract

Ascorbate (vitamin C) is an essential micronutrient in humans. The chronic severe deficiency of ascorbate, termed *scurvy*, has long been associated with increased susceptibility to infections. How ascorbate affects the immune system at the cellular and molecular levels remained unclear. Here, from a micronutrient screen, we identified ascorbate as a potent enhancer for antibody response by facilitating the IL-21/STAT3-dependent plasma cell differentiation in mouse and human B cells. The effect of ascorbate is unique, as other antioxidants failed to promote plasma cell differentiation. Ascorbate is critical during early B cell activation by poising the cells to plasma cell lineage without affecting the proximal IL-21/STAT3 signaling and the overall transcriptome. Consistent with its role as a cofactor for epigenetic enzymes, ascorbate potentiates plasma cell differentiation by remodeling the epigenome via TET (Ten Eleven Translocation), the enzymes responsible for DNA demethylation by oxidizing 5-methylcytosines into 5-hydroxymethylcytosine (5hmC). Genome-wide 5hmC profiling identified ascorbate responsive elements (E_AR_) at the *Prdm1* locus, including a distal element with a STAT3 motif overlapped with a CpG that was methylated and modified by TET in the presence of ascorbate. The results suggest that an adequate level of VC is required for antibody response and highlight how micronutrients regulate the activity of epigenetic enzymes to regulate gene expression. Our findings imply that epigenetic enzymes can function as sensors to gauge the availability of metabolites and influence cell fate decisions.

## Introduction

Vitamin C (VC) or ascorbate is an essential micronutrient for maintaining cell barrier integrity and protecting cells from oxidant damage^1,2^. Due to a mutation in *GULO* (L-gulono-gamma-lactone oxidase), humans are unable to synthesize ascorbate and have to depend on dietary absorbtion^3^. Long-term ascorbate deficiency causes a disease termed *scurvy*, which has been associated with weakened immune responses and higher susceptibility to pneumonia^4^. Although *scurvy* is rare nowadays, a survey in early 2000s estimated that the VC insufficiency rate is around 7.1% in the U.S^5^, and is even more prevalent in some sub-groups including the elderly, smokers, those with limited dietary intake and those with increased oxidative stress due to illnesses^6-9^). VC insufficiency may contribute to the variation of immune response against infections. Thus, it is critical to understand how VC regulates immune response at the cellular and molecular level.

In the past decades, researchers have explored using VC as a therapeutic treatment for diseases, such as infections and cancers, but the results have been mixed^10-17^. For instance, the studies of VC oral supplementation on the prevention of “common cold” or to enhance immune responses have been highly controversial, often confounded by different administrative routes and quantitative methods^18-21^. The results could potentially be confounded by factors including the pre-existing VC levels in the participants^22-25^; genetic variants in the specific VC transporters^7^; the ill-defined viral pathogens for “common cold”; and the evolution of viruses to infect humans regardless of VC status. Notably, excess oral supplementation does not increase the steady-state VC concentration beyond the physiological level maintained by absorption and excretion^18^. Recently, VC is injected intravenously to achieve a supraphysiologic concentration for treating sepsis and cancers^13,14,26^. The outcomes from these studies were ambiguous on whether the “mega dose” VC was effective in treating these diseases^10-13,26-28^. Nonetheless, VC supplementation via either route can address acute deficiency which may contribute to poorer outcomes^29-31^ and is likely to be necessary for certain conditions, especially for the population with VC insufficiency.

VC deficiency/insufficiency has been associated with dysregulated immune responses based on historical observation and experimental evidence. For instance, using *Gulo*-knockout mice that were unable to synthesize VC, studies have shown that VC deficiency negatively affects the response to influenza infection. One study showed that influenza infection increased the production of pro-inflammatory cytokines (TNF-α, IL-1β) and lung pathology without affecting viral clearance^32^. Another study showed that *Gulo*-deficient mice are defective in viral clearance due to decreased type I interferon response^33^. In addition to *Gulo*-deficient mice, guinea pigs are natural unable to synthesize VC and have been used as a model to show that VC is important for T-cell-dependent antibody response after immunization^34,35^. The mechanism by which VC regulates innate and adaptive immune responses remains unclear. In addition to its antioxidative function, VC is a cofactor for several epigenetic enzymes including Ten-eleven-translocation (TET)^36-38^. TET proteins (TET1, TET2, TET3) are critical for DNA demethylation by oxidizing 5-methylcytosine (5mC) mainly into 5-hydroxymethylcytosine (5hmC) and other oxidized bases, which are the intermediates for demethylation^39^. Besides enhancing neuronal and hematological differentiations *in vitro*^40^, VC facilitates the differentiation of regulatory T cells by potentiating TET-mediated demethylation of the *Foxp3* intronic enhancer *CNS2*^41-44^. These findings strongly suggested that one of the physiological functions of VC is to ensure the proper regulation of the epigenome that is required for coordinated immune responses.

B cells are responsible for the humoral immunity against pathogens and plasma cells are specialized in secreting antibodies. Plasma cells express the key transcription factor (TF) BLIMP1 (encoded by *Prdm1*) and can be induced by T-cell-derived signals, including CD40 ligand and IL-21. Using a two-step *in vitro* system modeling the T-dependent plasma cell differentiation, we performed a micronutrient screen and identified VC as a potent enhancer for plasma cell differentiation in B cells from mice and humans. The effect on plasma cells is specific to VC, as other antioxidants had no significant effect. We identified that the early B cell activation (1^st^ step) is the critical period requiring the presence of VC, which enables the B cells to become plasma cells after IL-21 stimulation. Intriguingly, VC treatment had only minimal influence on the transcriptome and IL-21 proximal signaling, suggesting the effect of VC is likely via the epigenome. Indeed, VC treatment increases the activity of TET as indicated by the level of 5hmC. Genome-wide 5hmC profiling of the *Prdm1* locus revealed several VC-responsive elements, including one of the elements (E58) with a highly conserved STAT3 motif overlapped with a CpG. The CpG at E58 is methylated in naïve B cells and potentially modified with 5hmC in the presence of VC. Using B cells deficient in *Tet2* and/or *Tet3*, we showed that TET2 or TET3 is sufficient to mediate the effect of VC. Our results suggest that an adequate level of VC is required for proper antibody response and highlight the influence of micronutrients on cell differentiation via epigenetic enzymes.

## Results

### Vitamin C as an Enhancer for Plasma Cell Differentiation

To study the role of micronutrients in plasma cell differentiation, we used a well-characterized B cell culture system^45^. In brief, mouse naïve B cells were cultured with 40LB, a stromal cell expressing CD40L and BAFF to activate and promote the survival of B cells (**Fig. 1A**). Naïve B cells were seeded with 40LB in the presence of IL-4 for four days (1^st^ step), followed by a secondary co-culture with IL-21 for three days to induce the differentiation of plasma cells (2^nd^ step) by the surface marker CD138 (Syndecan-1). Around 20% plasma cells were generated in this two-step culture after seven days (**Fig. 1B**).

**Figure 1.**
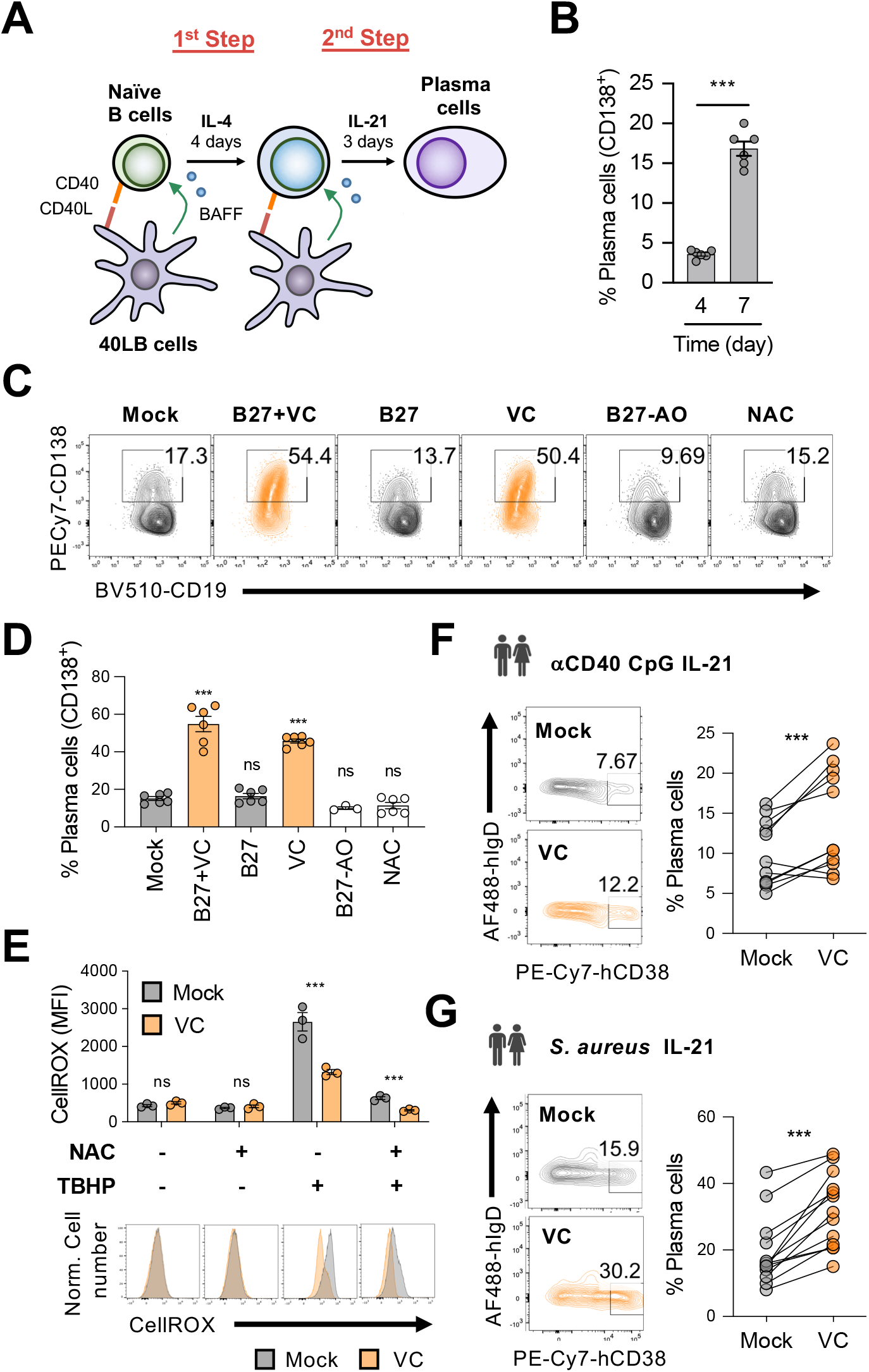
Vitamin C augments mouse and human plasma cell differentiation. (**A**) Schematic depiction of the 2-step 40LB cell culture system. Splenic naïve B cells were isolated and cultured with IL-4 and irradiated 40LB cells, which express CD40L and BAFF. Four days after culture, cells were sub-cultured to new layer of 40LB cells. IL-21 was used to induce plasma cell differentiation. (**B**) FACS analysis of plasma cells generated after each step during the 2-step culture. The plasma cell marker CD138, a plasma cell marker, was analyzed on day 4 and day 7 (n=6). (**C-D**) Micronutrient screen for plasma cell differentiation. B cells were cultured with supplements as indicated were analyzed by FACS on day 7 (n=6). (**E**) Low basal oxidative states in cultured B cells. B cells were cultured for 4 days with or without VC and the reactive oxygen species were monitored using a FACS-based fluorescent assay (CellROX). The antioxidant N-acetylcysteine (NAC) and oxidant tert-Butyl hydroperoxide (TBHP) were added 30 and 15 minutes, respectively, prior to the assay as controls (n=3). (**F**,**G**) VC enhances the differentiation of human plasma cells. Naïve B cells were isolated from human PBMCs and stimulated for 6 days as indicated to induced plasma cell differentiation in the presence or absence of VC. Data are from 5-6 independent experiments with 6 donors. All data are from at least two independent experiments. Mean ± SEM was shown for bar charts. Statistical significance was determined by unpaired (**B, E**) or paired (**F, G**) Student’s t test, and by one-way ANOVA with Dunnett’s post hoc test (**D**). ***, *P <* 0.001. *ns*, not significant.

We reasoned that the relatively inefficient plasma cell differentiation might be due to the paucity of micronutrients, as the deficiencies have been linked to increased susceptibility to infections^46^. To determine if effective B cell differentiation requires micronutrients, we included B27 and/or VC to the B cell culture. B27 is a culture supplement containing hormones, antioxidants (e.g. vitamin E and reduced glutathione), and other micronutrients (e.g. vitamins A, B7, selenium). The combination of B27 and VC was effective in promoting the differentiation of neurons^47-50^. Compared to control (Mock), the combination of B27 and VC significantly increased the percentage by at least 3-fold after the 7-day culture (**Fig. 1C, 1D**). Remarkably, VC was the major component responsible for the enhancement of plasma cell differentiation (**Fig 1C, 1D**), while B27 alone or B27 without antioxidants (B27-AO) have no effect. These culture supplements and VC had no significant effect on cell numbers and cell death (**Fig. S1A-C**), suggesting the increased plasma cells was not due to a preferentially increased survival.

Given that the primary activity of VC relies on its antioxidant potential, it is possible that VC relieves oxidative stress that may inhibit plasma cell differentiation. To address this possibility, we cultured B cells in the presence of another antioxidant N-acetylcysteine (NAC). Unlike VC, NAC had no significant effect on plasma cell differentiation (**Fig. 1C, 1D)**. In addition, B27 and the culture media also contained other antioxidants but were unable to promote plasma cell differentiation (**Fig 1C, 1D**), suggesting the effect on plasma cell is specific to VC and may not generalize to other antioxidants.

Next, we analyzed the oxidative state in the cultured B cells with a fluorescent dye (CellROX). The data showed that the mock and VC-treated cells had similarly low oxidative levels (left panel; **Fig. 1E**), which is supported by the fact that the addition of NAC during the CellROX assay did not significantly decrease the fluorescent signal (2^nd^ panel, **Fig. 1E**). To confirm the validity of the assay, tert-Butyl hydroperoxide (TBHP) was added to induce reactive oxygen species (ROS), which results in increased in the fluorescent signal (3^rd^ panel, **Fig. 1E**). The presence of VC in the culture media was sufficient to quench the exogenous ROS that can be further removed with the addition of NAC (3^rd^ and 4^th^ panels; **Fig. 1E**). The results demonstrated that the activated B cells in our system exhibited low oxidative stress, thus the effect of VC on plasma cell is likely independent of its general antioxidative function.

To address if activity of VC on plasma cells is conserved in human, we isolated naïve B cells from healthy donors and induced plasma cell differentiation using two conditions: 1) anti-CD40, toll-like receptor ligand CpG, and IL-21 in CpG (ODN); 2) *S. aureus* Cowan I (SAC) and IL-21. Consistent with its effect on mouse cells, VC can facilitate the differentiation of human plasma cells in both conditions (**Fig. 1F, 1G**). Together, the results suggested that VC is an important micronutrient for the generation of plasma cell in mice and humans.

### VC Supplement Facilitates the Differentiation of *bona fide* Plasma Cells

Our data revealed that VC promotes plasma cell differentiation indicated by using the surface expression of CD138. To confirm the results, we analyzed the protein expression of BLIMP1 and IRF4, two TFs important for plasma cell. Consistent with CD138, the percentage of BLIMP1 and IRF4 increased substantially after 7-day culture with VC (**Fig. 2A, 2B**). VC-treated B cells also downregulated the B-lineage TF PAX5 (**Fig. 2A upper panel; 2B**) similar to mature plasma cells. While B cells secreted minimal number of antibodies after the 1^st^ step culture regardless of VC (**Fig. S2A**), VC treatment significantly enhanced the antibody secretion after stimulation with IL-21 during 2^nd^ step culture (**Fig. 2C**). These results demonstrated that VC enhanced the differentiation of *bona fide* plasma cells.

**Figure 2.**
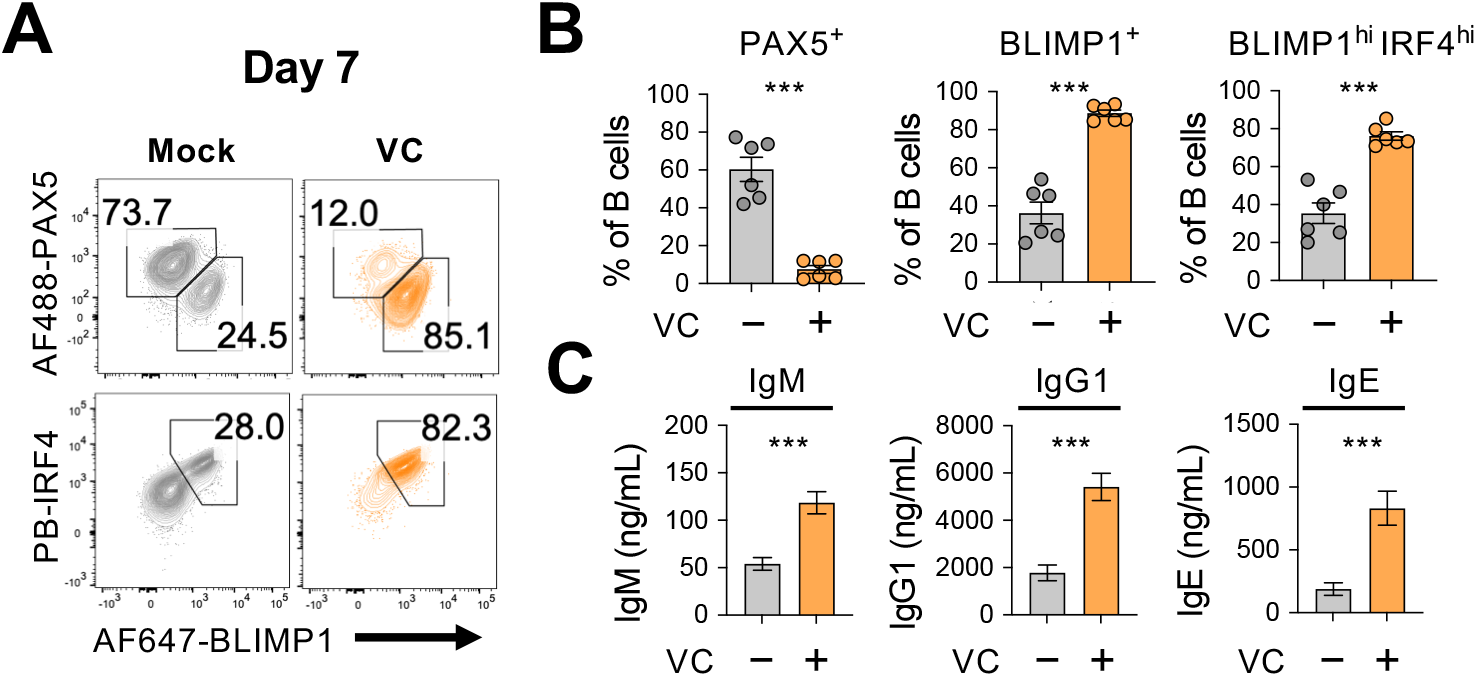
VC promotes the generation of *bona fide* plasma cells. (**A-B**) Expression analysis of lineage TFs. B cells were cultured as indicated for 7 days and the expression of transcription factors were analyzed by intracellular staining and FACS (n=6). Representative FACS plots **(A)** and summaries **(B)** are shown. Note that BLIMP1 and IRF4 are important TFs plasma cells differentiation. (**C**) Antibody secretion by VC-induced plasma cells. Culture supernatants were collected after day 7 and the secreted antibodies with indicated isotypes (IgM, IgG1, IgE) were analyzed by ELISA (n=6). All data are from at least two independent experiments. Mean ± SEM was shown for bar charts and the statistical significance was determined by unpaired Student’s t test. ***, *P* < 0.001.

### VC is Required for the Initial Activation Step of Plasma Cell Culture

As described above (**Fig. 1A**), the *in vitro* plasma cell differentiation is a two-step process, where naïve B cells are initially activated with IL-4 (1^st^ step) then stimulated with IL-21 to induce plasma cell differentiation (2^nd^ step). To identify the critical period for VC, we treated the cells with VC at different time points as indicated (**Fig. 3A**) and analyzed the cells by FACS on day 7 for CD138 expression. Consistent with the above result, the addition of VC throughout the culture significantly enhanced plasma cell differentiation compared to control (**Fig. 3A**; compare I vs IV). The data showed that the addition of VC during the 1^st^ step is sufficient to enhance plasma cell differentiation (**Fig. 3A; II**), while the effect is less pronounced when VC is added during the 2^nd^ step (**Fig. 3A; III**). The higher percentage of plasma cells after VC treatment is not due to increased kinetics of plasma cell differentiation. B cells were similar in function and phenotype after the initial four days of culture regardless of VC treatment (**Fig. 3B, S2A, S2B**). These data demonstrated that VC is important for conditioning the B cells during the initial activation period.

**Figure 3.**
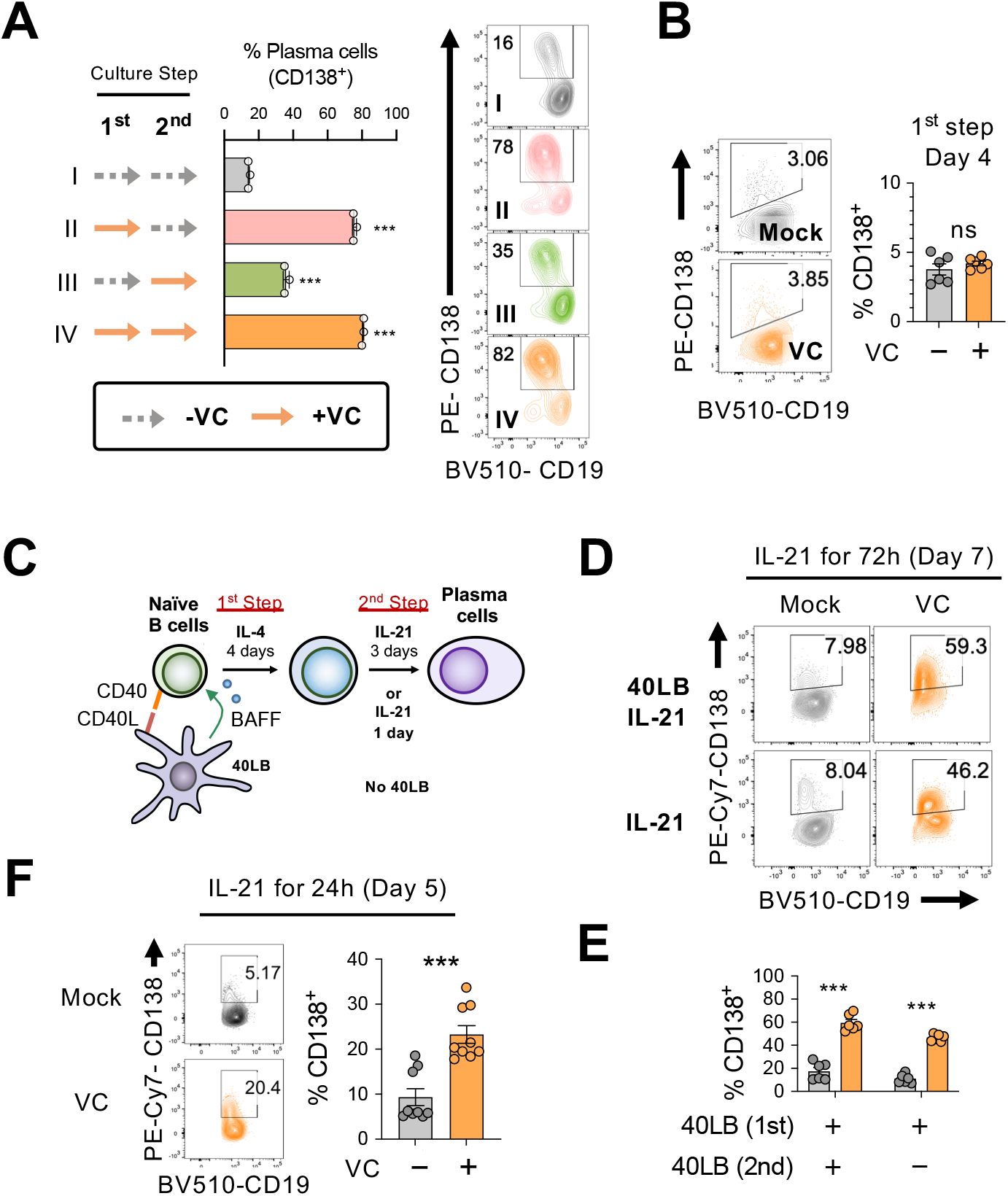
VC potentiates the IL-21-dependent differentiation of plasma cell during the initial activation phase. (**A**) VC is required during activation phase. B cells were treated with or without VC as indicated for the 1^st^ and 2^nd^ step culture. Dotted arrow represents control and solid orange arrow indicates VC. Plasma cell differentiation (%CD138^+^) was analyzed on day 7 by FACS. Representative FACS plots were shown on right and summarized data are shown on left. Roman numbers are used for correlation between panels. (**B**) VC has no significant effect on the percentage of plasma cells after 1^st^ step culture. Representative FACS plots for plasma cell marker CD138 of B cells on day 4 (left) and the summarized data (right) are shown (n=6). (**C-E**) IL-21 alone is sufficient to induce plasma cell differentiation during 2^nd^ step. (**C**) Schematic representation of the B cell culture used in the next panels. B cells were cultured with IL-21 for indicated time without the 40LB stromal cells. (**D-E**) Cells were either cultured with or without 40LB in the presence of IL-21 during the 2^nd^ step. Percentage of plasma cells (%CD138^+^) was analyzed by FACS on day 7 (n=6). The representative FACS plots (**D**) and summarized data (**E**) are shown. (**F**) IL-21 stimulation induced plasma cell differentiation after 24h. B cells were culture in the absence (mock) or presence (VC) of VC during the 1^st^ step with IL-4 and 40LB. Cells were stimulated with IL-21 alone for 24h and the percentage of CD138^+^ cells was analyzed by FACS (n=9). All data are from at least two independent experiments. Mean ± SEM was shown for bar charts and the statistical significance was determined by one-way ANOVA with Dunnett’s post hoc test (**A**) and unpaired Student’s t test (**B, E, F**). ***, *P* < 0.001. ns, not significant.

During the 2^nd^ step culture, B cells are stimulated with 40LB and IL-21 to induce plasma cell differentiation. To determine whether VC facilitates plasma cell differentiation by modulating the CD40L/BAFF or cytokine signals, we cultured the B cells with IL-21 in the presence or absence of 40LB cells during the 2^nd^ step as indicated (**Fig. 3C**). The results show that VC significantly enhances plasma cell differentiation regardless of 40LB cells by measuring CD138 (**Fig. 3D-E**), antibody secretion (**Fig. S3A**), and TF expression (**Fig. S3B**). While the percentage of plasma cells was similar between mock- and VC-treated cells after 1^st^ step culture (**Fig. 3B**), stimulation with IL-21 for 24h was sufficient to induce significantly more plasma cells from VC-treated B cells (**Fig. 3F**). These results suggest that VC conditions B cells to be responsive to the IL-21-induced plasma cell differentiation.

### VC has Limited Effect on the Proximal IL-21 Signaling and Transcriptome

IL-21 is produced by T cells and can induce the phosphorylation of STAT3 after binding to IL-21 receptor (IL-21R) on B cells. To address if VC promotes plasma cell differentiation by altering IL-21 signaling, we cultured the B cells with 40LB and IL-4 for four days, and the expression of IL-21R was analyzed by flow cytometry. The data showed that IL-21R expression was comparable between mock- and VC-treated cells (**Fig. 4A**). To test if VC may enhance the signaling capacity of IL-21R, we stimulated the cells cultured the cells with IL-21 for 30 minutes and the phosphorylation of STAT3 (pSTAT3) was analyzed. Similarly, the results showed that VC has no significant effect on the phosphorylation of STAT3 in response to IL-21 (**Fig. 4B**). Note that the same concentration of IL-21 was used for both the short-term stimulation (30 minutes) and the 2^nd^ step culture. These data suggest that VC-mediated conditioning of the B cells is not due to heightened responsiveness to IL-21.

**Figure 4.**
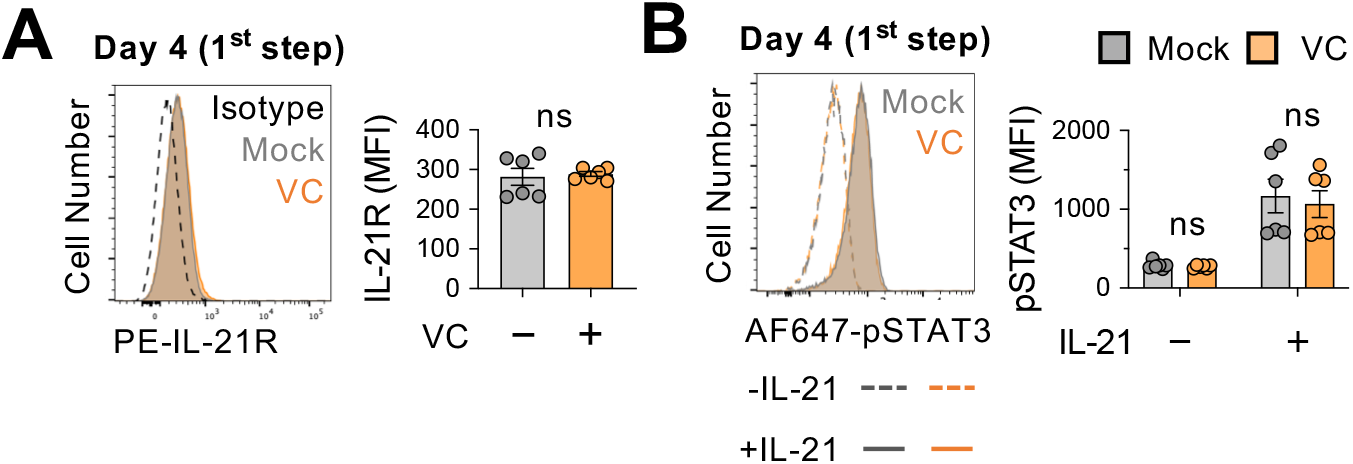
VC did not promote the proximal IL-21 signaling. **(A) VC has no effect on IL-21 receptor expression**. Mock or VC-treated B cells were cultured for 4 days and the expression of IL-21R was analyzed by FACS (n=6). **(B) VC does not alter the IL-21 induced STAT3 phosphorylation**. B cells from day 4 culture were stimulated with IL-21 for 30 minutes and the level of STAT3 phosphorylation (pSTAT3) was analyzed by FACS (n=6). Representative histograms are shown in left and summarized data in right. All data are from at least two independent experiments. Mean ± SEM was shown for bar charts and the statistical significance was determined by unpaired Student’s t test. ns, not significant.

Based on the above results, VC poises B cells toward plasma cell lineage without affecting the overall phenotypes (**Fig. 3B, S2**) and IL-21 signaling (**Fig. 4**) prior to IL-21 stimulation. To identify the potential genes regulated by VC, we analyzed the transcriptomes of naïve and B cells cultured for four days in the presence of absence of VC by RNA-seq. As expected, B cell activation induced substantial changes in transcriptomes, with thousands of differentially expressed genes (DEGs; **Fig. 5A, 5B**). Surprisingly, the transcriptomes of B cells are highly similar between control and VC-treated cells (**Fig. 5C**), with only one and five genes differentially decreased and increased, respectively. Among the DEGs, the majority of them are non-coding transcripts (e.g. *Gm48840, Gm13192, Gm5340*) and proteins with unknown function (1700120K04Rik, *Lrp2bp*). One of the genes induced by VC is Rnf165/Ark2C, a ubiquitin E3 ligase that has been associated with impaired motor neuron axon growth^51^. Its function in the immune system remained unclear. However, based on the minimal number of DEGs, we speculated that VC might condition the B cells without affecting the transcriptome.

**Figure 5.**
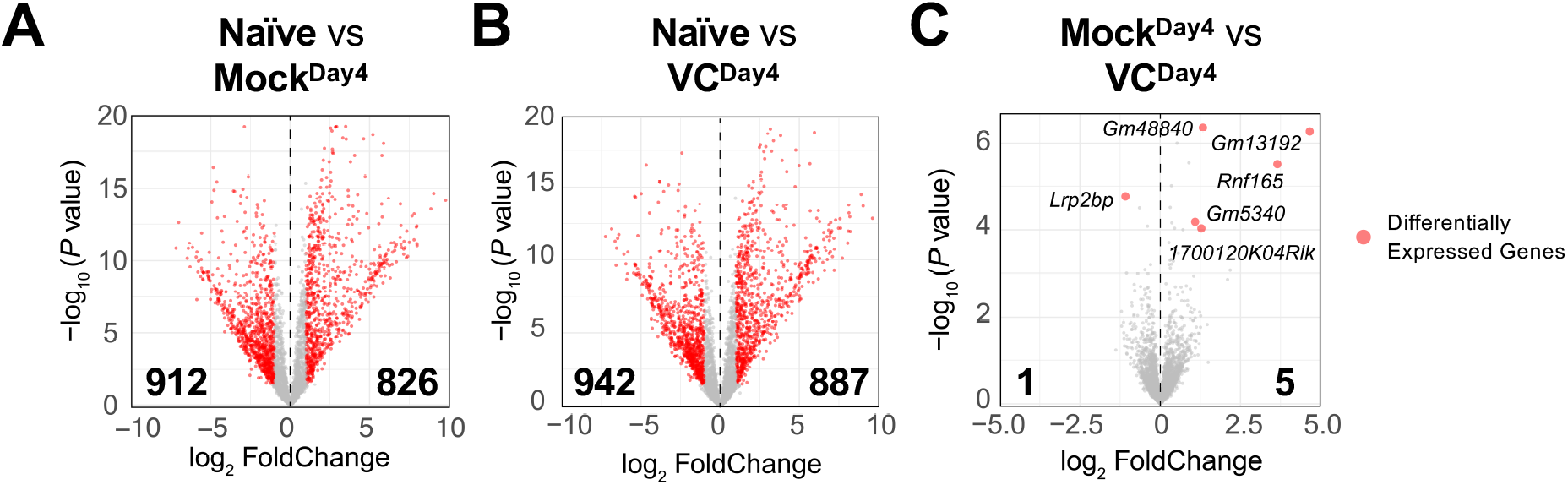
Limited effect of vitamin C on the transcriptome in activated B cells. The mRNA was isolated from naïve and indicated day 4 activated B cells and the transcriptomes were analyzed by RNA-seq (see Materials and Methods). Comparisons shown are between (**A**) Naïve vs mock; (**B**) Naïve vs VC; (**C**) Mock vs VC. Three independent biological replicates were used for each group. Red dots indicate differentially expressed genes with an adjusted *P* value ≤ 0.01 and log_2_ fold change ≥ 1. Numbers indicate the numbers of differentially expressed genes. Gene names for all differentially expressed genes are shown on (**C**).

### The Effect of VC is Dependent on either TET2 or TET3

In addition to being an general antioxidant, VC has an unique property to reduce Fe(III) to Fe(II) for epigenetic enzymes, including Ten-Eleven Translocation (TET). family proteins (TET1, TET2, TET3) are essential for DNA demethylation by oxidizing 5-methylcytosine (5mC) to mainly 5-hydroxymethylcytosine (5hmC)^39,40,52^. Previous studies have shown that TET2 and TET3 are required for B cell development and function^53-59^. Therefore, we hypothesize that VC may affect the B cell epigenome by enhancing the enzymatic activity of TET. We isolated B cells from control (*Cd19*^*Cre/+*^), *Tet2-*deficient (*Cd19*^*Cre/+*^ *Tet2*^*fl/fl*^), or *Tet3-*deficient (*Cd19*^*Cre/+*^ *Tet3*^*fl/fl*^) mice and cultured them in the presence or absence of VC. While VC greatly increased the percentage of plasma cells generated in control B cells (**Fig. 6A-D;** WT), the differentiation was slightly impaired in the *Tet2-* or *Tet3-*deficient cells (**Fig. 6A-D**). As TET2 and TET3 often function redundantly, we isolated B cells from *Tet2 and Tet3* conditional double-deficient mice to address if the effect of VC is mediated by TET enzymes. Since the deletion of *Tet2* and *Tet3* using *Cd19*^*Cre*^ resulted in aggressive B cell lymphoma at an early age (unpublished observation), we used a tamoxifen-inducible *Tet2/3-*deletion system to circumvent this caveat as previously described^56^. In this system, *Cre*^*ERT2*^ *Tet2*^*fl/fl*^ *Tet3*^*fl/fl*^ and control *Cre*^*ERT2*^ mice were injected with tamoxifen for five consecutive days to induce the deletions of *Tet2* and *Tet3*. B cells were isolated on day eight and cultured as above. Remarkably, the effect of VC on plasma cell differentiation dramatically decreased in the absence of TET2 and TET3 as measured by percentages of CD138^+^ (**Fig. 6E and 6G)** or BLIMP1^+^ (**Fig. 6F and 6H**) B cells. Therefore, these results strongly suggested that the ability of VC to enhance plasma cell differentiation is via TET2 and TET3.

**Figure 6.**
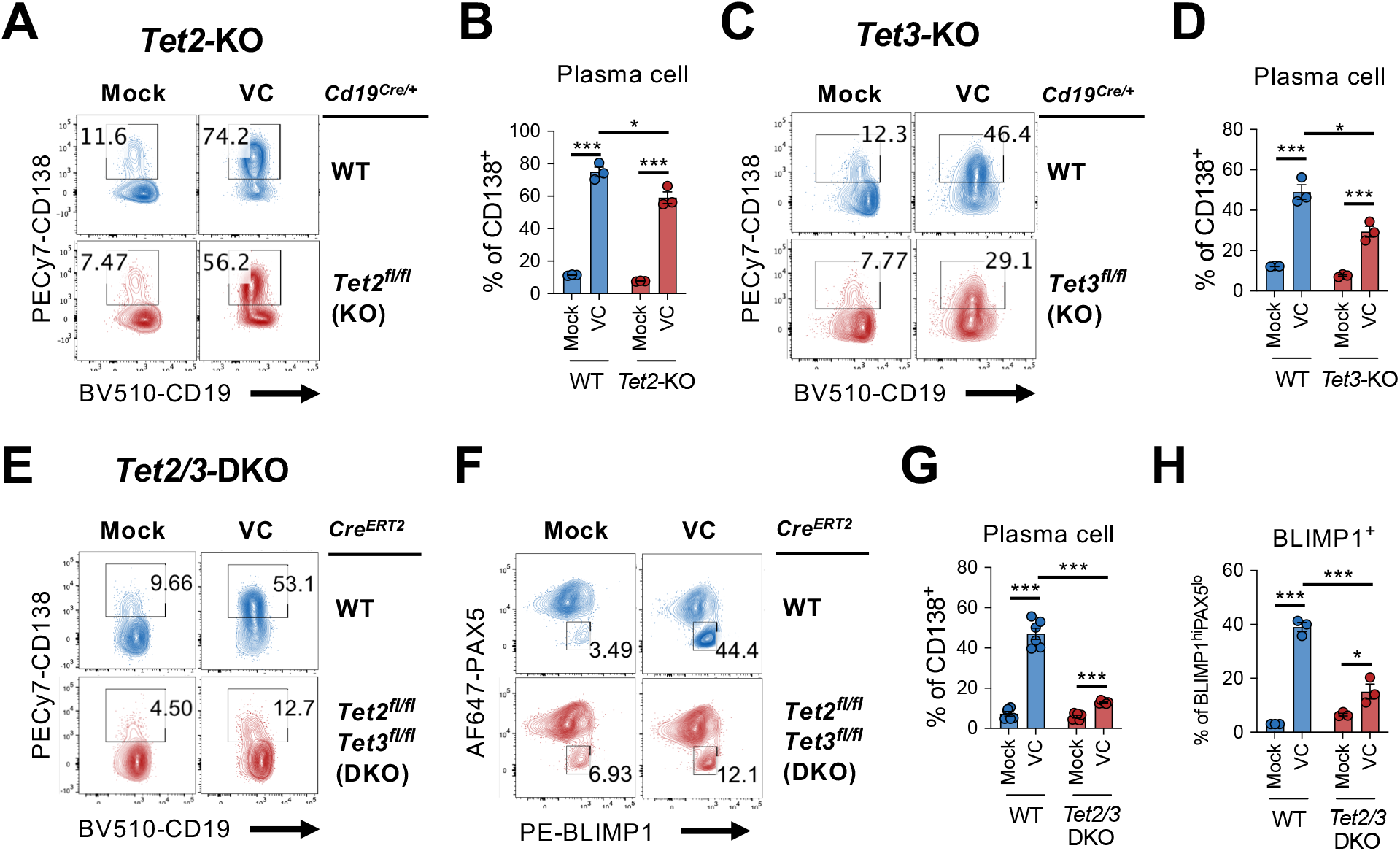
TET2 or TET3 is required for VC-mediated enhancement of plasma cell differentiation. (**A-D**) *Tet2* or *Tet3* is sufficient for VC-mediated increased in plasma cells. (**A-B**) B cells from WT (*Cd19*^*Cre/+*^) or *Tet2* conditional deficient (*Cd19*^*Cre/+*^ *Tet2*^*fl/*fl^; *Tet2*-KO) mice isolated and cultured as in Fig. 1A with or without VC. Percentage of CD138^+^ cells was analyzed by FACS on day 7 (n=3). (**C-D**) B cells were isolated from either WT (*Cd19*^*Cre/+*^) or *Tet3* conditional KO (*Cd19*^*Cre/+*^ *Tet3*^*fl/fl*^; *Tet3*-KO) and cultured as in (**A**; n=3). Representative FACS plots (**A, C**) and summarized data (**B, D**) are shown. (**E-H**) *Tet2* or *Tet3* are required for the effect of VC. Control (*Cre*^*ERT2*^) or *Tet2/Tet3* conditional deficient (*Cre*^*ERT2*^ *Tet2*^*fl/*fl^ *Tet3*^*fl/fl*^) mice were injected with tamoxifen for 5 consecutive days and B cells were isolated by cell sorting on day 8. The expression of *Rosa26-* YFP was used as a surrogate marker for Cre activity. B cells were cultured as in (**A**) or (**C**) and percentage of CD138 (**E**) and intracellular staining of TFs (**F**) were analyzed by FACS on day 7. Summarized data are shown in (**G**) and (**H**). Representative FACS plots for plasma cell marker CD138 (**E**) and transcription factors (**F**) on day 7. All data are from at least two independent experiments. Mean ± SEM was shown for bar charts and the statistical significance was determined by unpaired Student’s t test. ***, *P* < 0.001. *, *P* < 0.05. ns, not significant.

### VC Remodels the Genome-wide 5hmC Modification

TET enzymes oxidize 5mC into oxidized bases including 5hmC, a stable epigenetic medication and an intermediate for DNA demethylation^40,52,60^. To confirm VC indeed enhances TET activity, we used DNA dot blot to semi-quantitatively measure the level of 5hmC in B cells. Naïve B cells had the highest density of 5hmC compared with B cells from day four or day seven culture (**Fig. 7A**), an observation consistent with the passive dilution of 5hmC after cell divisions. The addition of VC significantly increased the amount of 5hmC compared to those in the mock control B cells at both time points (**Fig. 7A**; compare Mock and VC). These results confirm that VC enhances TET activity in B cells.

**Figure 7.**
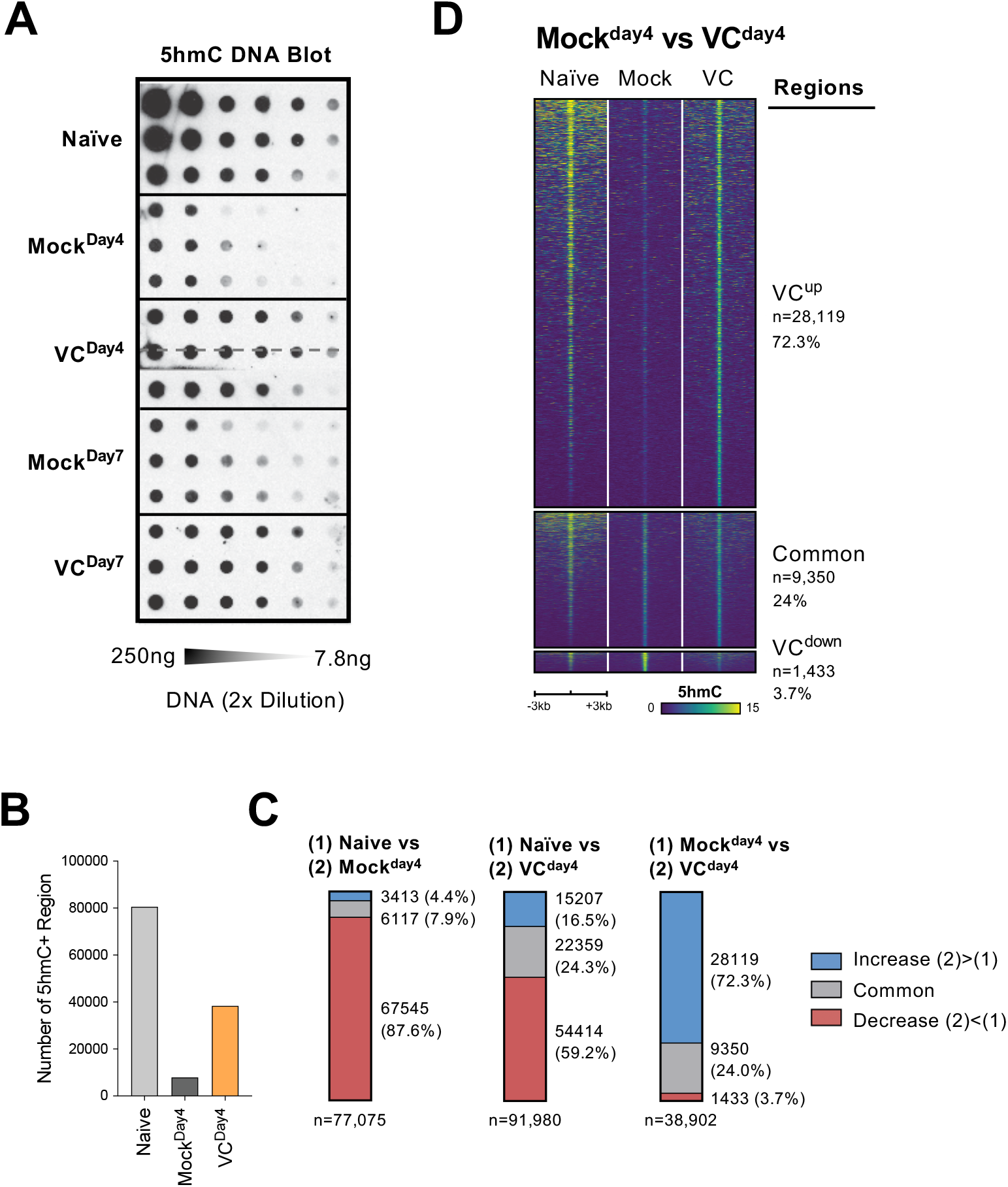
VC induced a global increase in 5hmC modification. (**A**). VC increased total 5hmC in B cells. DNA from indicated B cells were isolated and the total 5hmC levels were analyzed using cytosine 5-methylenesulphonate (CMS) dot blot (Materials and Methods). Note that CMS is the immunogenic product of 5hmC after bisulfite treatment. DNA was serially 2-fold diluted starting from 250ng to ∼7.8ng. Three biological replicates were blotted for each group. (**B-D**) The genome-wide 5hmC enrichment was analyzed using a CLICK-chemistry based pulldown method (HMCP). Two biological replicates were used for each condition (naïve, Mock^Day4^, VC^Day4^; see Materials and Methods for details). (**B**) VC treatment increased the numbers of 5hmC-enriched region. The number of 5hmC-enriched (5hmC+) regions were called using HOMER and the numbers of consensus regions between replicates are plotted. (**C**) VC maintains pre-existing and induces *de novo* 5hmC-enriched regions. Pairwise comparisons between the three groups are shown. Numbers and percentages of regions are shown for each comparison. Colors indicate the 5hmC status of the regions. Number of 5hmC+ region in naïve B cells and B cells with or without VC treatment. (**D**) Visualization of 5hmC at differential regions between B cells from Mock^Day4^ and VC^Day4^ groups. The 5hmC enrichment was plotted as heatmaps around (+/-3kb) the differential and common regions between Mock^Day4^ and VC^Day4^ (right panel of **7C**). Normalized 5hmC counts are plotted as color scale and each row represent a region. **Figure 7-source data 1**. Original blot for Fig. 7A CMS dot blot.

To identify the targeted genomic regions regulated by TET, we used HMCP, a modified CLICK-chemistry-based 5hmC pulldown method, to analyze the genome-wide 5hmC distribution. Overall, the number of 5hmC-enriched regions correlated with the levels of 5hmC (**Fig. 7A**): naïve B cells contained the most 5hmC-enriched regions; B cells activated for four days (Mock^Day4^) had the lowest; B cells cultured with VC (VC^Day4^) had significantly more 5hmC-enriched regions compared to mock control (**Fig. 7B**). Analysis of the differentially hydroxymethylated regions (DhmRs) revealed that B cells cultured for four days (Mock^Day4^) lost the majority (87.6%; **Fig. 7C** left panel and **Fig. S4A**) of the 5hmC-enriched regions that were identified in naïve B cells. Smaller percentages of peaks are either maintained (7.9%) or gained (4.4%). In contrast, while considerable numbers of DhmRs were decreased in B cells cultured with VC (VC^Day4^; 59.2%; **Fig. 7C** middle panel and **Fig. S4B**), a significant number of regions either maintained (24.3%) or gained 5hmC (16.5%) after culture. Most of these 5hmC-enriched regions were distal to the transcription start sites (**Fig. S4C-E**), suggesting that many of these regions might be potentially regulatory elements as previously described^56^.

To understand the effect of VC on TET-mediated epigenome remodeling, we focused on the regions between the activated B cells from mock and VC groups. The analysis of DhmRs revealed that VC induced a significant number of 5hmC-enriched regions (72.3%; **Fig. 7C** right panel and **Fig. 7D**), while a small number of regions have decreased 5hmC (3.7%). Motif analysis of the regions enriched in VC-treated cells showed modest enrichment of motifs from TF families, including basic helix-loop-helix (bHLH), nuclear receptor (NR), and zinc fingers (Zf; **Fig. S5A**). The DhmRs preferentially found in mock control were enriched in bHLH, basic leucine zipper (bZIP), and Zf family TFs (**Fig. S5B**). How and if these TFs may collaborate with TET to facilitate DNA modification remains to be addressed. Altogether, the data demonstrated that VC enhanced the enzymatic activity of TET, resulting in a substantial increase in 5hmC throughout the genome.

### Potential *Tet*-responsive element at the *Prdm1* distal region

*Prdm1-*encoded BLIMP1 is the TF critical for the differentiation and function of plasma cells. The above results showed that VC promotes plasma cell differentiation via TET-mediated deposition of 5hmC. Therefore, we hypothesize that VC may affect the DNA modification status at the *Prdm1* locus, resulting in an increased permissiveness to IL-21-mediated upregulation. In naïve B cells, the *Prdm1* gene body and several downstream regions, including a previously described 3’ *Prdm1* enhancer close to the transcriptional terminal site, are enriched in 5hmC (**Fig. 8A**, top panel). In B cells activated without VC (Mock^Day4^), the majority of the regions showed decreased 5hmC. The decrease was more noticeable after considered the lower global 5hmC (**Fig. 7A**). Remarkably, VC induced the enrichment of 5hmC at least 14 locations (teal bars in Fig. 8B, *MockDay4 vs VCDay4*). One of the most pronounced enrichment was at E58, a previously undescribed element located at +58kb relative to the start of *Prdm1*^61^. DhmR analyses showed that E58 is preferentially enriched in VC-treated B cells compared to both naïve and mock control (**Fig. 7B**). Moreover, overlaying a previously published DNA methylation data demonstrated that E58 is methylated in naïve B cells and is at the edge of a demethylating region (**Fig. 7C**), consistent with a previous observation where the boundaries between methylated and demethylation regions are enriched in 5hmC^62^.

**Figure 8.**
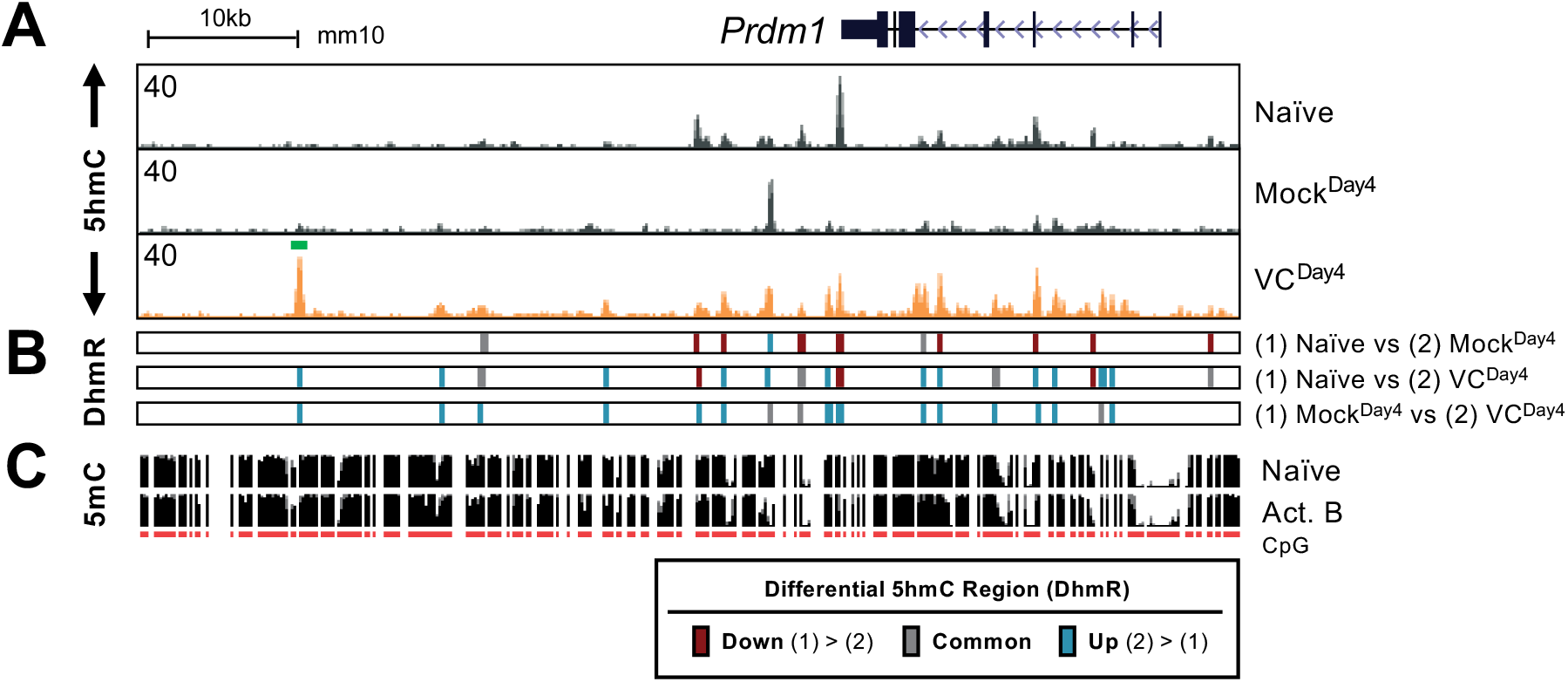
VC induced 5hmC modification at the *Prdm1* locus. (**A**) The genome browser tracks of the 5hmC enrichment at the *Prdm1* locus. Data shown are the average of two biological replicates as in **Fig. 7D. (B)** Differential 5hmC-enriched regions (DhmRs) between the indicated group as in **Fig. 7C**. Colors indicate the differential status of the regions as depicted in the legend. **(C)** DNA methylation in naïve and 48h-activated B cells from previous publication^56,72^. The height of the black bars indicated the percentage of CpG methylation and the red track (CpG) indicated the CpGs that were covered in the analysis. Note that VC induced several 5hmC-enriched regions when compared to control (teal lines in 3^rd^ row of DhmR) and we termed these regions E_AR_ (ascorbate response elements). One of the prominent E_AR_ is E58 (labeled in green in **A**), which showed significant increase in 5hmC after VC treatment.

Since VC induced 5hmC modification at several locations at the *Prdm1* locus, we speculate that some of these *cis*-elements may be responsible for the effect of VC on plasma cells (**Fig. 3D, 3E, S3B**). Among the elements, E58 has one of the highest proportions of 5hmC modification at the *Prdm1* locus. Coincidentally, analysis of previously published STAT3 ChIP-seq in T cells^61^ showed that IL-21-induced STAT3 could bind to E58 (**Fig. S6A**). Interestingly, a predicted STAT3 motif at E58 contained a CpG site that is methylated in naïve B cells (**Fig. S6B**). This CpG-containing STAT3 motif is highly conserved (**Fig. S6C**), suggesting that the motif may have significant biological functions and is potentially regulated by DNA modification. In conclusion, our data showed that plasma cell differentiation is sensitive to the level of VC, which can affect the epigenome by modulating the activity of epigenetic enzymes including TET. Heighten TET activity may increase the probability of the DNA modification/demethylation at critical regulatory elements and may skew the lineage decision of the differentiating B cells.

## Discussion

VC has long been known for its broad effect on immune responses. Long-term VC deficiency results in *scurvy*, a disease historically associated with increased susceptibility to infections^4,14,63^. While recent studies have focused on VC supplements on disease treatments or prevention^10,12-14,20,28,30,31,63^, relatively little is known about the mechanism of how VC deficiency may affect the immune response.

Using an *in vitro* 2-step model of plasma cell differentiation, we performed a limited micronutrient screen and identified VC as a strong potentiator of plasma cell differentiation in mouse and human B cells (**Fig. 1C-D, 1F-G**). The effect of VC on plasma cells is likely independent of its general antioxidant capacity. First, other antioxidants, including vitamin E, reduced glutathione, NAC, and 2-mercaptoethanol (a component in the complete media), could not recapitulate the effect (**Fig. 1C-D**). Second, the basal levels of oxidative stress were low in the cells regardless of VC treatment (**Fig. 1E**), suggesting that VC may exert its activity on other cellular processes. We further identified that the main effect of VC occurred during the initial activation step (1^st^ step; **Fig. 3A**), prior to the stimulation with IL-21. Intriguingly, after the four days of 1^st^ step culture, the percentage of plasma cells (**Fig. S2, 3B**), the capacity of proximal IL-21/STAT3 signaling (**Fig. 4**), and the transcriptomes (**Fig. 5C**) were remarkably similar between control and VC-treated cells. As recent studies suggested that VC is a cofactor for epigenetic enzymes, we speculated that VC might facilitate B cells to differentiate into plasma cells by remodeling the epigenetic landscape (**Fig. S7**).

Our data showed that VC indeed has a profound effect on the epigenome, including the genome-wide distribution of 5hmC. VC increased the total 5hmC deposited by TET (**Fig. 7A**), and either TET2 or TET3 is required for VC-enhanced plasma cell differentiation (**Fig. 6**). The genome-wide mapping of 5hmC showed that in the absence of VC, the majority of 5hmC-enriched regions in naïve B cells showed decreased 5hmC after B cell proliferated (Naïve vs Mock^Day4^; **Fig. 7B, 7C**). While B cells cultured with VC showed a similar decrease in the numbers of 5hmC-enriched regions (naïve vs VC^Day4^; **Fig. 7B, 7C**), VC was able to promote TET activity and resulted in the maintenance of a subset of original 5hmC-enriched regions. Moreover, VC was required for the appearance of at least 10,000 *de novo* 5hmC peaks induced by activation (3,413 in Mock^Day4^ vs 15,207 in VC^Day4^; **Fig. 7C**). The data suggest that TET2 or TET3 may be recruited to these regions in the activated B cells but the enzymatic rate is limited when the intracellular level of VC is low.

Since *Prdm1*/BLIMP1 is the TF essential for plasma cell differentiation, we speculate that VC may affect the 5hmC modification at the *Prdm1* locus and render it permissive to STAT3-mediated upregulation. Upon examination of the 5hmC at the *Prdm1* locus, we identified at least 14 VC-induced DhmRs in the B cells cultured with VC (VC^Day4^) that we termed “ascorbate responsive elements” (E_AR_; **Fig. 8B**), which are defined as the regions with gained 5hmC after VC treatment. One noticeable E_AR_ is E58, an undescribed *cis-*element located at +58 kb, and several known enhancers^61^. Previously published STAT3 ChIP-seq data showed that STAT3 could bind to E58 in response to IL-21 in T cells (**Fig. S6A**)^61^, coinciding with a conserved STAT3 binding motif at E58 (**Fig. S6B, S6C**). Interestingly, the STAT3 motif contains a CpG site overlapping the 5hmC signal (**Fig. 8A, 8C**). Based on this observation, we suggest that VC alters the DNA modification status at E58 and potentially other E_AR_ at the *Prdm1* locus that may affect the recruitment of TFs. The mechanism behind VC-mediated plasma cell differentiation remained to be addressed.

Our data is accordant with a recent publication showing that VC can promote plasma cell differentiation via TET enzymes^64^. Similar to our results, Qi et al. showed that the effect of VC on plasma cells is dependent on TET, and TET-deficient affects VC-assisted DNA demethylation at the *Prdm1* intronic enhancers. While the previous study provided *in vivo* data, we have shown detailed molecular mechanisms of how VC enhances plasma cell differentiation. For instance, we dissected the 2-step culture system and have shown VC is critical during the initial B cell activation (1^st^ step). The effect of VC is unique and cannot be substituted by other antioxidants. Moreover, we demonstrated that VC has no immediate effect on the transcriptome and IL-21/STAT3 signaling after the 1^st^ step culture. Our data demonstrated that the main effect of VC is to remodel the epigenome. Furthermore, our genome-wide profiling of 5hmC revealed several E_AR_ at the *Prdm1* locus, including a potential novel element E58 that might be sensitive to the methylation status. Together, the studies here and Qi et al. have presented a clear picture of the importance of VC on regulating B cell differentiation.

Increasing evidence has revealed the link between the metabolome and epigenome in the immune system^65^. For instance, VC stabilizes the *in vitro* induced regulatory T cells by stabilizing the expression of lineage TF *Foxp3* by facilitating TET-mediated DNA demethylation at CNS2, a *Foxp3* intronic enhancer^41,43,44^. Another example is α-ketoglutarate (αKG), an intermediate in the Kreb cycle and a substrate for many epigenetic enzymes. Studies have shown that αKG modulates macrophage function by facilitating histone demethylation^66^ and promoting IL-2-sensitive gene expression in T cells by TET-mediated DNA demethylation^67^. Our results and others have shown that VC is a vital micronutrient required for efficient plasma cell differentiation^64^. Based on these observations, we hypothesize that epigenetic enzymes may act as rheostats to control appropriate gene transcription by integrating the availability of micronutrients or metabolites. Future studies will be needed to address how micronutrients, including VC, contribute to immune response via epigenetic remodeling.

## Acknowledgements

Drs. Anjana Rao, Patrick Hogan (LJI), Andrew McKnight, Rachael Soloff, Anne-Laure Perraud (Kyowa Kirin Pharmaceutical Research; KKR) for the supports and advice; Roberta Nowak and Samantha Blake for assistant (LJI); Cheryl Kim and Denise Hinz at for cell sorting; the Institute for Genomic Medicine at Nationwide Children’s Hospital for NGS-sequencing; Drs. Eugene Oltz and Ken Oestreich (OSU) for advice and critical reading; Dr. Hazem Ghoneim for assistant with experiments; Drs. Emily Hemann, Gang Xin, and Adriana Forero (OSU) for providing critical reagents during the peak of the pandemic; and all the Lio lab members for their contribution and critical reading of the manuscript. 40LB cells were a gift from Dr. Daisuke Kitamura (Tokyo University of Science). This research was funded by LJI/KKR young investigator fund; NIH National Cancer Institute K22 (K22CA241290); startup funds from the Department of Microbial Infection and Immunity and from the Pelotonia Institute of Immuno-oncology at the Ohio State University (all to C.-W.J.L.).

## Materials and Methods

### Animal Experiments

All animal experiments were approved by IACUC at La Jolla Institute and Ohio State University. C57BL/6J mice (6-12 weeks; Jax#000664) were purchased from Jackson Laboratory (Bar Harbor, ME). *Tet2*^*fl/fl*^ *Tet3*^*fl/fl*^ (generous gifts from Dr. Anjana Rao) crossed with *Rosa26*^*LSL-EYFP*^ (Jax#006148) and *Ubc-Cre*^*ERT2*^ (Jax#008085; *Cre*^*ERT2*^ herein) mice were as described previously^56^. To induce *Tet2/3*-deletion with Cre^ERT2^, mice were injected with 2mg of tamoxifen in corn oil (Millipore-Sigma, St. Louis, MO) for 5 consecutive days and rested for 2 days before the isolation of primary B cell.

### Cell culture

40LB cells were cultured in D10 media consists of DMEM (high-glucose), 10% fetal bovine serum (FBS; Gemini Bio, West Sacramento, CA), and 2mM GlutaMax. Primary B cells were cultured in R10 media consists of RPMI1640, 10% FBS (Gemini Bio), 2mM GlutaMax, 1x non-essential amino acid, 1mM sodium pyruvate, 10mM HEPES, 50 ug/mL gentamicin, and 55 uM 2-mercaptoethanol. B27 serum-free supplement (50x) was added as 1x. The components of B27 are listed in **Table S4**. All cell culture reagents above are from Gibco-ThermoFisher (Waltham, MA) unless otherwise stated.

### Induced germinal center (iGB or 40LB) culture

The iGB culture was performed according to previous publication with slight modifications^3^, Briefly, 40LB cells were irradiated (30 Gy) and 1×10^5^ cells were plated for each well in a 24-well tissue-culture plate in D10 media. The following day, B cells were isolated from mouse spleens using EasySep mouse B cell isolation kit (Stemcell Technologies, Vancouver, Canada) according to manufacturer’s instruction. For the first step culture, B cells (10,000) were plated with the irradiated 40LB cells in R10 media with 1 ng/mL recombinant mouse interleukin-4 (rmIL-4; Peprotech, East Windsor, NJ). On day 4, B cells were resuspended and 50,000 cells were transferred to irradiated 40LB cells in R10 media with 10 ng/mL rmIL-21 (Peprotech) for the second step culture. Cells were analyzed on day 7. For VC treatment, L-ascorbic acid 2-phosphate (Millipore-Sigma) was dissolved in water and sterilized with 0.22um syringe-filter and was added at a final concentration of 100 µg/mL (310.5 µM). VC was added on the beginning of the 1^st^ and 2^nd^ steps co-culture.

### Isolation and culture of primary human B cells

Human peripheral blood mononuclear cells (PBMCs) were isolated from healthy donors with IRB approval (to D.J.W.). Briefly, 20 to 40ml of whole blood was diluted with phosphate based-saline (PBS) and overlaid on Ficoll-Paque Plus (Cytiva, Marlborough, MA). After centrifugation, PBMCs were transferred from the interface and washed with PBS. Primary human B cells were isolated from PBMCs using the EasySep Human Naïve B cell isolation kit (Stemcell Technologies). Naïve B cells were cultured with or without VC (100 µg/mL) in R10 media. To induce plasma cell differentiation, cells were cultured with recombinant human IL-21 (100 ng/mL; Peprotech) and either with: 0.01% PANSORBIN (Millipore); or ODN (2.5 µM; Invivogen, San Diego, CA) and anti-hCD40 (1 µg/mL; BioLegend). Cells were harvested on day six for flow cytometry analysis.

### Flow Cytometry

Cells were resuspended in FACS buffer (1% bovine serum albumin, 1mM Ethylenediaminetetraacetic acid/EDTA, 0.05% sodium azide) and incubated with Fc-receptor blocking antibody 2.4G2 (5 µg/mL; BioXCell, Lebanon, NH) for at least 10 mins on ice. Fluorescence-conjugated antibodies and live-dead dye were added and incubate on ice for 25-30 mins. Cells were washed at least two times with FACS buffer, fixed with 1% paraformaldehyde (ThermoFisher), and analyzed by either FACS Canto-II, LSR-II, or Accuri C6 (all BD, Franklin Lakes, NJ). FACS Aria II (BD) and MA900 (Sony Biotechnology, San Jose, CA) were used for cell isolation. For intracellular transcription factor staining, cells were stained by using True-Nuclear transcription factor buffer set (Biolegend, San Diego, CA). Briefly, cells were fixed with 1x fix concentrate buffer (Biolegend) at room temperature (RT) for 45-60 mins and washed twice with 1x permeabilization buffer. Cells were resuspended in 1x permeabilization buffer and stained with fluorescence-conjugated antibodies at RT for 30 mins. Cells were washed twice with 1x permeabilization buffer and prior to FACS analysis. For detection of apoptosis, 1×10^6^ cells were washed with PBS and resuspended in 1x binding buffer (10 mM HEPES pH 7.4, 140 mM NaCl, 2.5 mM CaCl_2_). Biotin-conjugated Annexin V (Stemcell Technologies) was added and incubated at RT for 10-15 mins. Cells were washed and resuspended in 1x binding buffer. APC-Streptavidin (Biolegend) was added and incubated at RT for 10-15 mins. Cells were washed, pelleted, and resuspended in 1X binding buffer. 7-Aminoactinomycin D (7-AAD; BD) was added to label the dead cells by incubating at RT for at least 5 mins and analyzed by FACS immediately. FACS antibodies and reagents are listed on Table S5.

### STAT3 Phosphorylation Assay

Cells from the 1^st^ 40LB culture were washed once with complete media and stimulated with rmIL-21 (10 ng/mL final) at a concentration of 5×10^5^ cells/mL at 37ºC for 30 mins. Cells were fixed immediately with paraformaldehyde (final 2%; ThermoFisher) and incubated at 4ºC for 30 mins. Cells were washed twice with FACS buffer and the surface antigens were stained with fluorescence-conjugated antibodies at RT for 30 mins. Cells were washed twice again with FACS buffer and fixed with ice-cold methanol (final 90%) and stored at -20ºC overnight. The next day, cells were washed at least twice with FACS buffer to remove the fixative before incubating with the Fc-receptor blocking antibody 2.4G2 (BioXCell) and rat serum (Stemcell Technologies), followed by the anti-phosphoSTAT3 (Tyr705) antibody (Biolegend) at RT for 30 mins. Cells were washed twice with FACS buffer, fixed with 1% paraformaldehyde (ThermoFisher), and analyzed by FACS.

### Cytosine-5-methylenesulfonate (CMS) dot blot

CMS dot blot was performed as previously described^2^. Briefly, genomic DNA was treated with Methylcode bisulfite conversion kit (ThermoFisher) to convert 5hmC into CMS. DNA was diluted, denatured, neutralized, and immobilized on a nitrocellulose membrane with the Bio-Dot apparatus (Bio-Rad, Hercules, CA). CMS was detected using rabbit anti-CMS antisera (gift from Dr. Anjana Rao) followed by peroxidase-conjugated goat-anti-rabbit IgG secondary antibody. Membrane was exposed using SuperSignal West Femto (ThermoFisher) and X-ray films.

### 5hmC enrichment analysis

Genomic DNA was isolated from B cells either using Blood and Tissue or Flexigene kits (Qiagen, Hilden, Germany). DNA was quantified with Qubit (ThermoFisher) and sonicated to around 150-200bp using Bioruptor Pico (Diagenode, Denville, NJ). 5hmC enrichment analysis was performed using the HMCP kit (Cambridge Epigenetix, Cambridge, United Kingdom) based on the addition of azido-glucose to 5hmC by T4-beta-glucosyltransferase following the manufacturer’s instruction. The 5hmC-enriched and input libraries were sequenced using Novaseq 6000 (Illumina, San Diego, CA) with 50×50bp paired-end using SP flow cells (Nationwide Children Hospital, Columbus, OH).

### RNA sequencing

RNA was purified using RNeasy Kit (Qiagen) and was analyzed by TapeStation RNA tape (Agilent, Santa Clara, CA). Total RNA (RIN>9.5) was used for mRNA isolation using the NEBNext Poly(A) mRNA Magnetic isolation module (NEB, Ipswich, MA). Libraries were constructed using NEBNext Ultra II Directional RNA Library Prep Kit (NEB) according to the manufacturer’s protocol and barcoded using the NEBNext Multiplex (unique dual index) Oligos (NEB). Libraries were sequenced using Novaseq 6000 (Illumina) with 50×50 bp paired-end using SP flow cells (Nationwide Children Hospital).

### ELISA

Culture supernatants were collected on day four and day seven of 40LB culture and stored at -20C. Capture antibodies for specific isotypes (goat anti-IgM, -IgG, and IgE; Southern Biotech, Birmingham, AL) were coated on high-binding 96-well plates (Corning, Tweksbury, MA) at 1 µg/mL overnight. After blocking with 1% bovine serum albumin in PBS, standards (IgM and IgG from Southern Biotech; IgE from Biolegend) and samples were applied to the coated plate for 1 hr at room temperature (RT). After washed with PBS-T (PBS with 0.05% Tween-20), 160 ng/ml of HRP-conjugated donkey anti-mouse IgG (H+L) detection antibody (Jackson Immunoresearch, West Grove, PA) was added to the plates and incubate for 1 hr at RT. The peroxidase substrate tetramethylbenzidine (TMB; ThermoFisher) was added and the reactions were stopped by adding 50 uL of 1 M sulfuric acid. The absorbance (OD) of the plates were read using SpectraMax (Molecular Device, San Jose, CA) at 450nm (using OD at 540nm as background). Antibody concentrations were extrapolated from the standards.

### Oxidative stress analysis

The ROS levels were detected using CellROX deep red flow cytometry assay kit (ThermoFisher) according to the manufacturer’s protocol. Briefly, 50,000 cells cultured for four days with 40LB were added to a 24-well plate. Cells might be treated with 250 uM NAC for 1hr at 37°C. Tert-butyl hydroperoxide (TBHP; 200 μM), an inducer of ROS, might be added to the cells for 30 mins 37°C. To monitor the oxidative stress, 2uL of CellROX Deep Red reagent was added to the cells for 15 mins followed by 1uL of SYTOX Blue Dead Cell stain solution for another 15 mins. The cells were analyzed immediately by flow cytometry.

### Cell sorting

*Cre*^*ERT2*^ *Tet2*^*fl/fl*^ *Tet3*^*fl/fl*^ *Rosa26*^*LSL-EYFP*^ mice and *Cre*^*ERT2*^ *Rosa26*^*LSL-EYFP*^ (control) were injected intraperitoneally with tamoxifen (Sigma; 2mg per mouse in corn oil) for 5 consecutive days and rested for 2 days before the isolation of primary B cell. CD19^+^ YFP^+^ live splenic B cells were sorted using MA900 with 100µm flow chips (Sony Biotechnology).

### Bioinformatics analyses

#### 5hmC analysis

Paired-end reads were mapped to the mouse genome mm10 GRCm38 (Dec. 2011) from UCSC using Bowtie 2 (v2.4.2)^68^, reads that failed the alignment and duplicates were removed using Samtools (v1.10). Transcripts mapping to autosomal and sex chromosomes were kept. Bigwig files were generated using bamCoverage (--normalizeUsing RPKM) using Deeptools (v3.5.1)^69^. Peaks were called using MACS2^70^ with control samples (callpeak -g 1.87e9 -q 0.01 -- keep-dup all --nomodel –broad). Reads were compared to the blacklisted regions for mm10 (ENCODE 2016). Count matrix was generated using DiffBind (v2.0.2)^71^ dba count with normalization DBA_SCORE_TMM_MINUS_EFFECTIVE. Differentially enriched 5hmC regions were determined using edgeR, regions were selected by an adjusted p-value lower than 0.05. Heatmaps were generated using Deeptools.

#### RNA-seq analysis

Paired-end reads were mapped to the mouse genome using GRCm38 genome sequence, primary assembly (ENCODE 2016) using STAR (v 2.7.0). Gene count matrix was generated with STAR using Comprehensive gene annotation CHR Regions. Unmapped regions were filtered. Differential gene expression was calculated using limma (v3.12) using voom transformation. Genes were selected by an adjusted p-value (FDR) lower than 0.05 and a log(2) fold enrichment higher or equal to 1.

#### Published STAT3 ChIP-seq

The Bigwig files for STAT3 ChIP-seq were downloaded from Cistrome (CistromeDB # 4577 and 4580). The data were from CD4 T cells stimulated with IL-21 (100 ng/mL) for 1 hour^61^.

#### Data Availability

Codes used in this study will be available upon requested. The data have been deposited to National Center for Biotechnology Information Gene Expression Omnibus (GEO GSE183681).

## Figure Legends

**Supplementary Figure 1.**
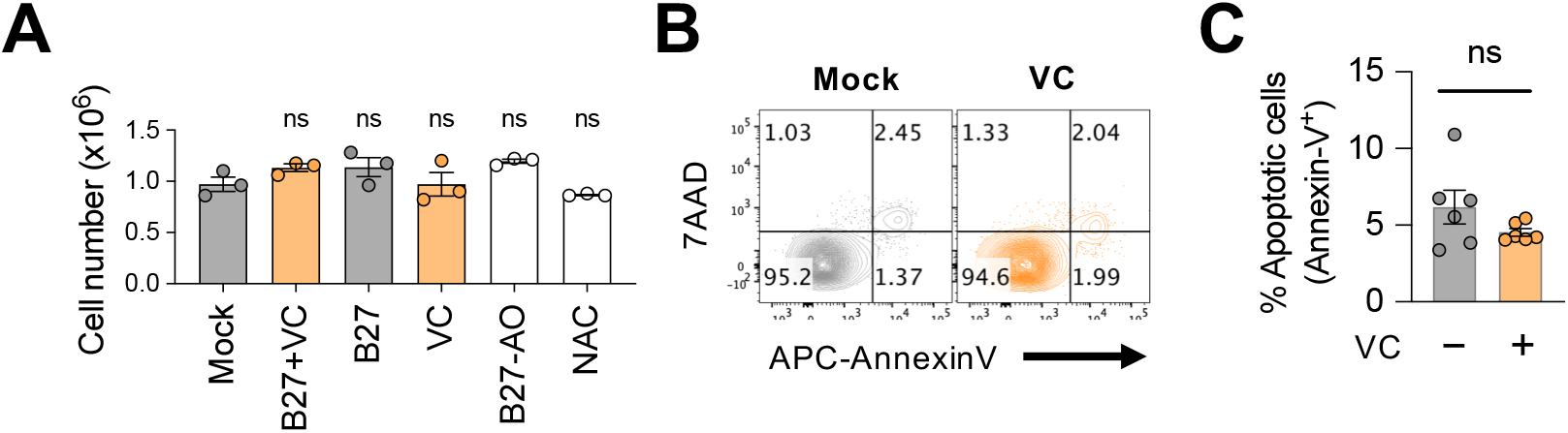
Minimal effect of VC on B cell survival. **(A)** Culture supplements had no significant effect on B cell numbers. B cells were cultured as in Fig. 1C-D and the cells were enumerated on day 7 using ViCell automated cell counter (n=3). **(B-C)** VC had no major effect on cell death. B cells were cultured with or without VC and the percentage of apoptotic cells were analyzed by FACS using 7AAD and AnnexinV (n=6). (B) Representative FACS plots. **(C)** Quantification. Statistical significance was determined by one-way ANOVA with Dunnett’s post hoc test **(A)** or unpaired Student’s t test **(C)**. ns, not significant (*P > 0*.*05*).

**Supplementary Figure 2.**
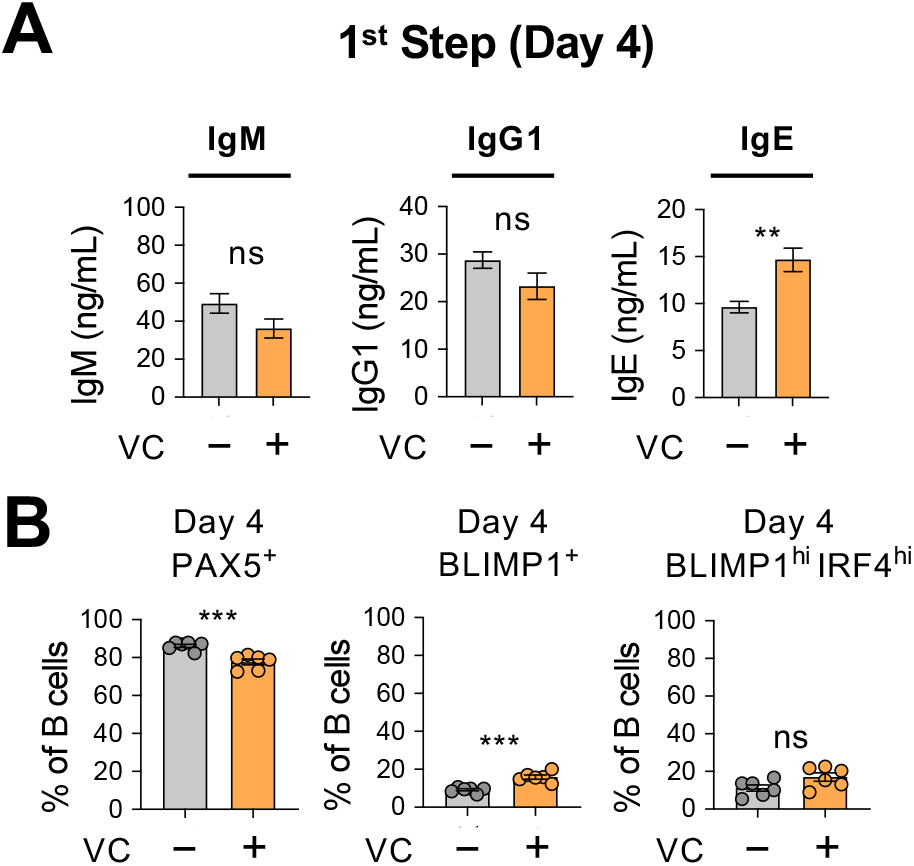
Minimal antibody secretion and differentiation of plasma cells at early stage. **(A)** Low level of antibody secretion on day 4. Mock or VC-treated B cells were cultured with 40LB and IL-4 for 4 days (1^st^ step; Fig. 1A) and the antibody secretion was analyzed by ELISA as in Fig. S1 (n=6). Note that the scales of IgG1 and IgE on Fig. 2C are at least 50-200x higher. (**B**) Low percentage of plasma cells on day 4. TFs were analyzed as in Fig. 2A-B. There are statistically significant but minor changes (compared Fig. 2B) in PAX5^+^ and BLIMP1^+^ cells between mock and VC groups (n=6). All data are from at least two independent experiments. Mean ± SEM was shown for bar charts and the statistical significance was determined by unpaired Student’s t test. ***, *P* < 0.001. **, *P* < 0.01.

**Supplementary Figure 3.**
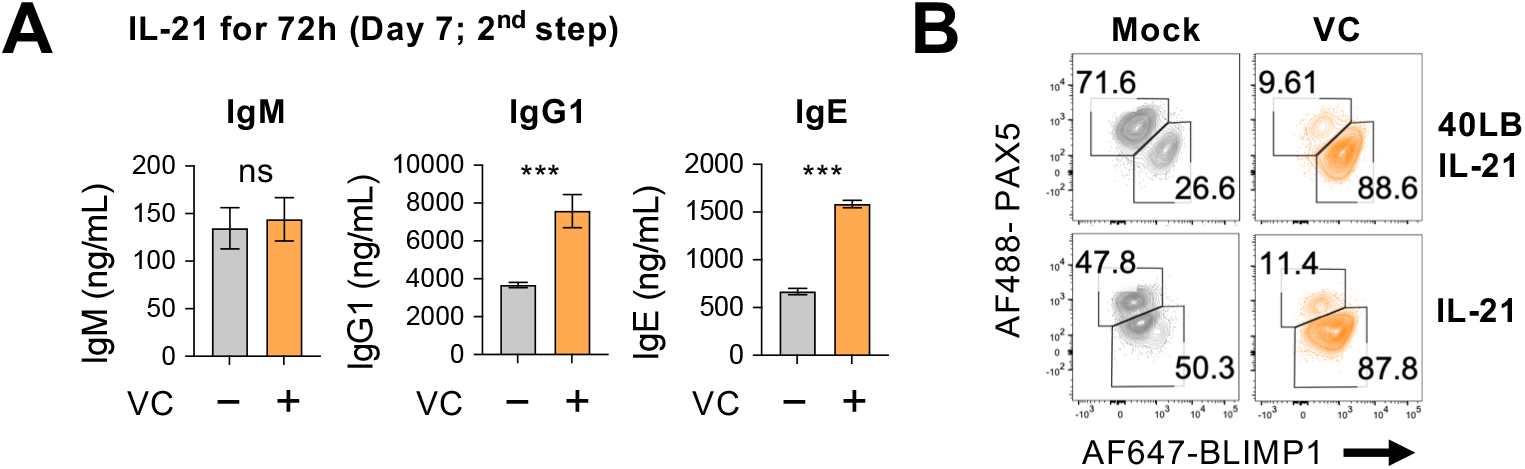
IL-21 alone during later phase is sufficient for plasma cell differentiation. B cells were cultured with IL-21 alone with or without 40LB during the 2^nd^ step as in **Fig. 3D. (A)** Antibodies secretion was analyzed by ELISA and **(B)** the expression of TFs was analyzed by FACS on day 7 (n=6). All data are from at least two independent experiments. Mean ± SEM was shown for bar charts and the statistical significance was determined by unpaired Student’s t test. ***, *P* < 0.001. ns, not significant.

**Supplementary Figure 4.**
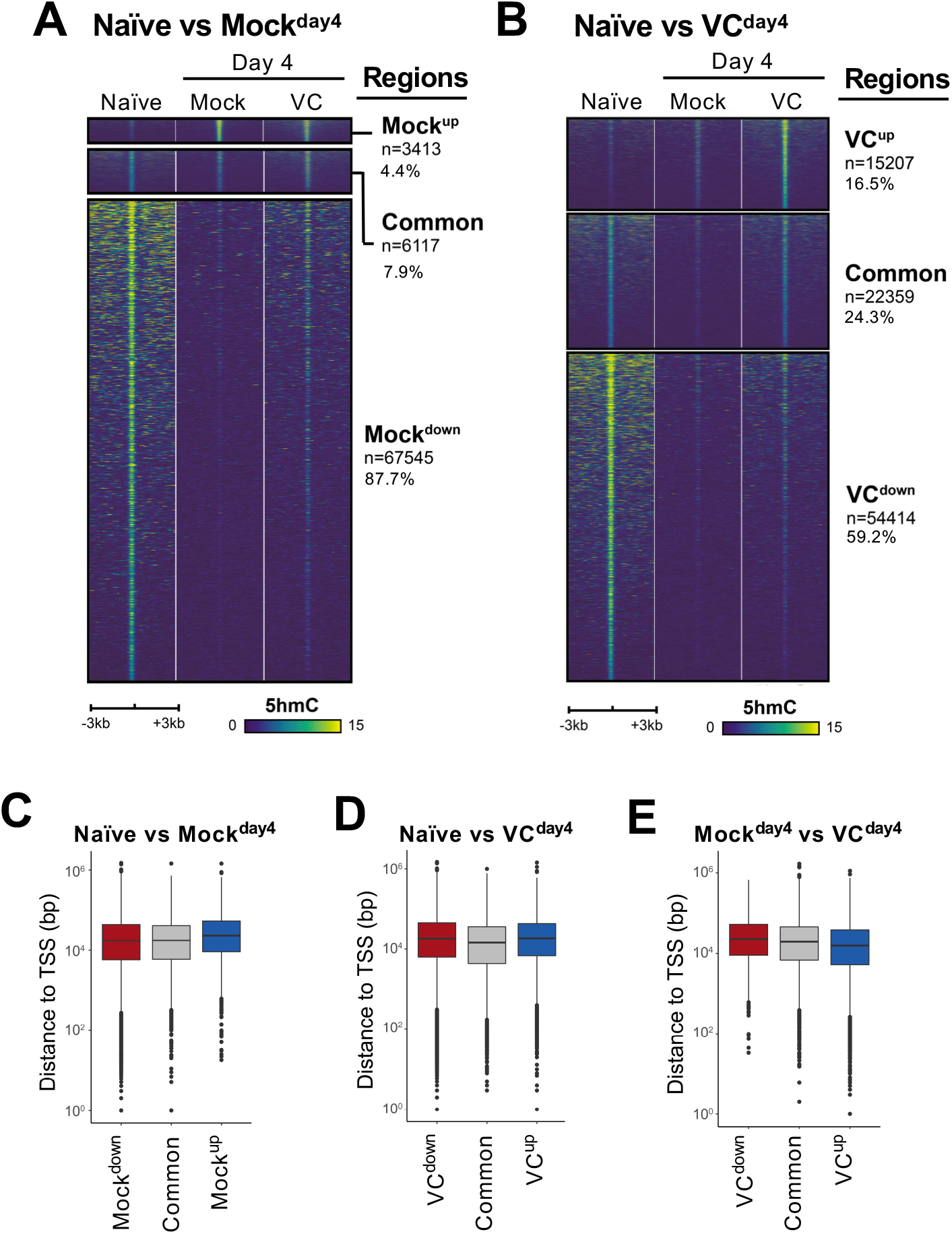
Analyses of 5hmC-enriched regions. **(A-B)** Heatmaps showing 5hmC enrichment at the differential and common regions between (A) Naïve and Mock^Day4^; (B) Naïve and VC^Day4^. Data were plotted as described in **Fig. 7D. (C-E)** Majority of 5hmC-enriched regions are distal elements. The distance between the indicated 5hmC-regions and the closest transcriptional start sites (TSSs) are plotted. Note that majority of the regions are at least 10 kb away from TSSs.

**Supplementary Figure 5.**
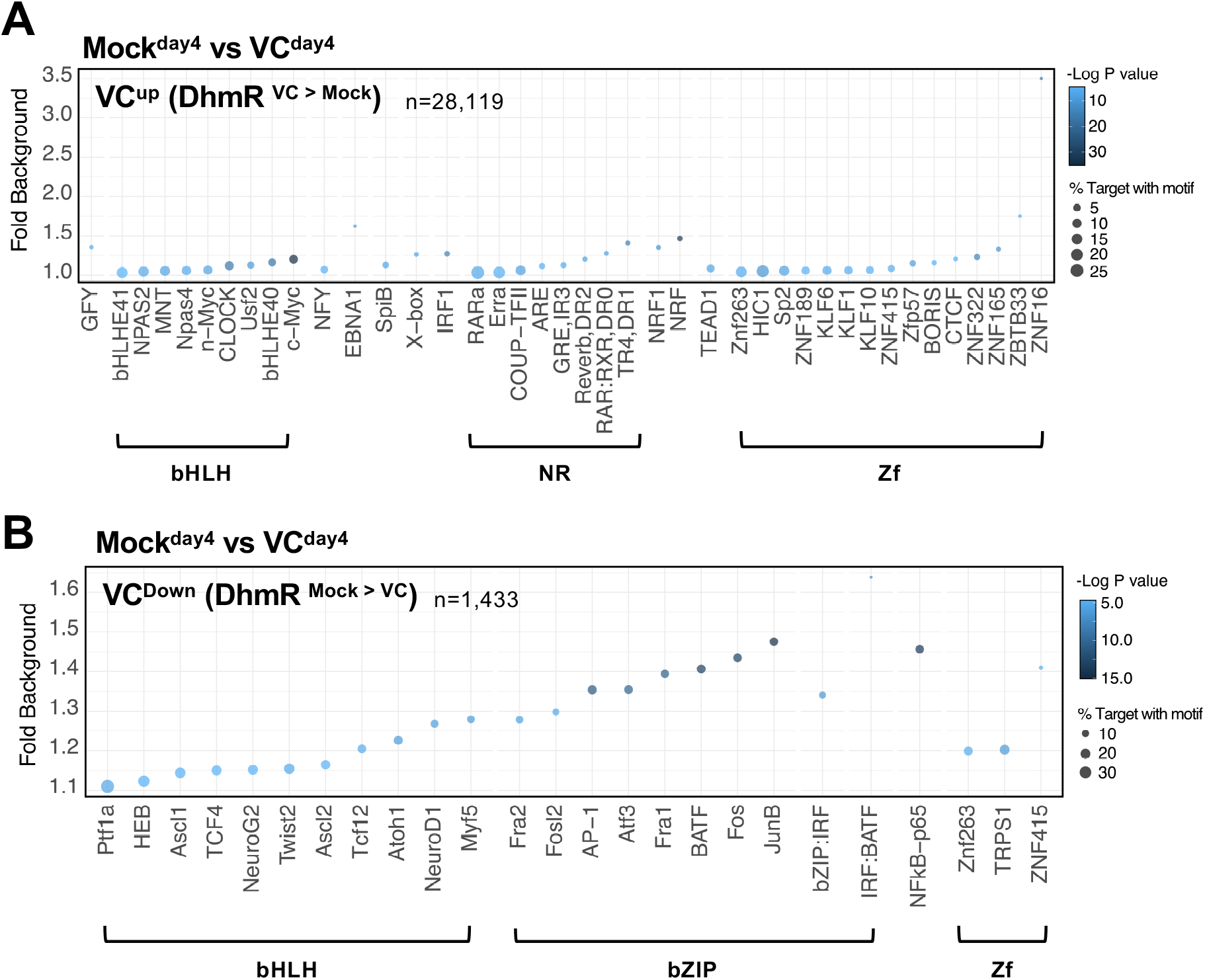
Enrichment of TF motifs at differential 5hmC regions. The enrichment of motifs was analyzed for the differential regions between Mock^Day4^ and VC^Day4^ using HOMER. Common regions between the two groups are used as backgrounds for motif analysis. Motif enrichment for regions with **(A)** increased in 5hmC (VC^UP^) and **(B)** decreased in 5hmC (VC^Down^) in the presence of VC. Y axis indicates the relative number (fold) of TF motifs in the differential regions compared to background regions. The size of the bubble indicates the percentage of all regions with the indicated TF motif (bottom). The bubble color indicates the statistical significance.

**Supplement Figure 6.**
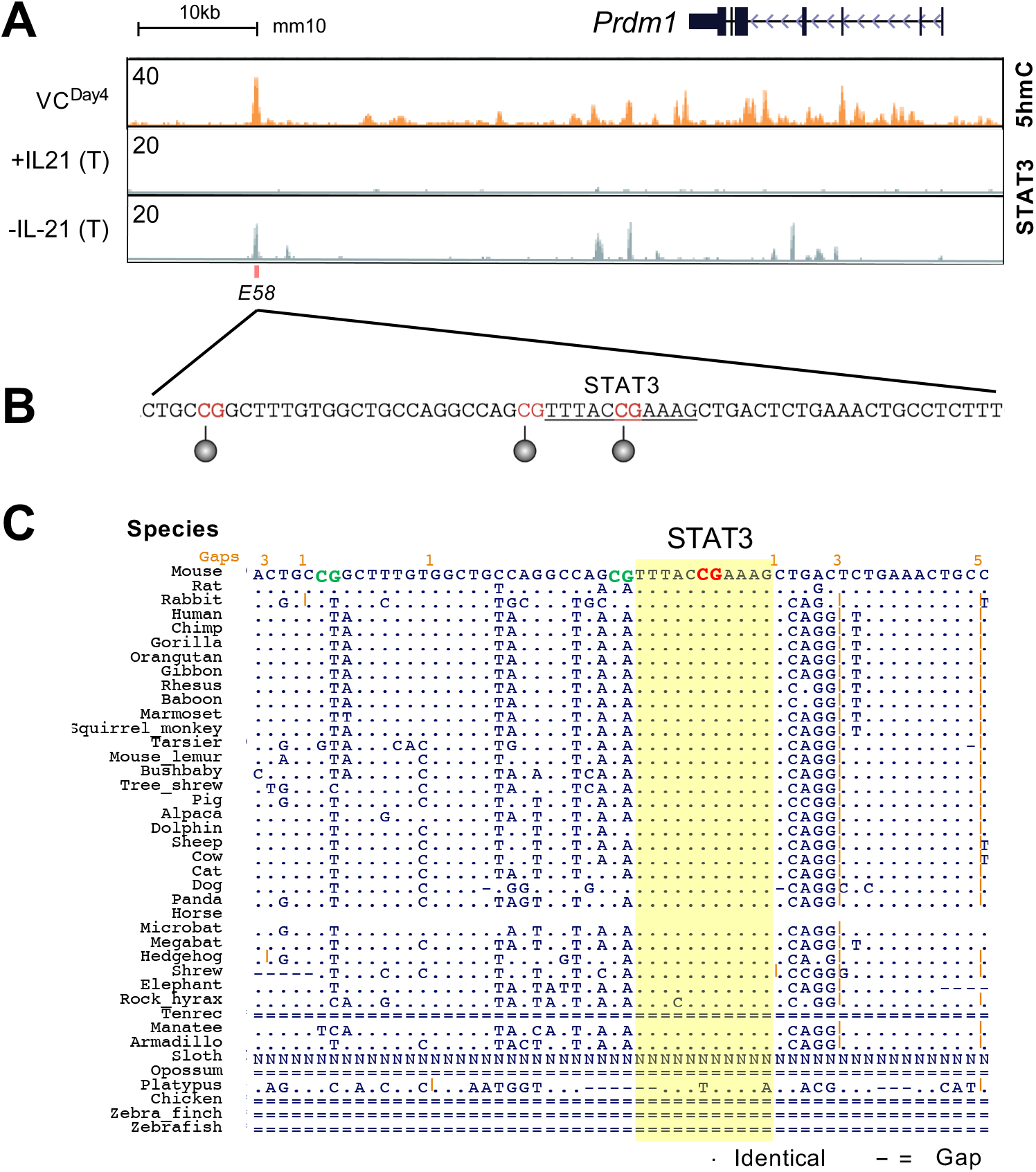
Sequence analysis of a potential *Prdm1* enhancer E58. **(A)** Genomebrowser tracks showing the 5hmC enrichment at the *Prdm1* locus from VC^Day4^ group (as in **Fig. 8A**). The previous published STAT3 ChIP-seq tracks from CD4 T cells stimulated with or without IL-21 were shown^61^. E58, one of the E_AR_, is depicted. **(B)** A potential STAT3 motif at E58 overlapped with a CpG that is methylated in naïve B cells (**Fig. 8C**). Motif analysis was performed using JASPAR 2020. **(C)** The CpG-containing STAT3 motif is highly conserved. The DNA sequences alignment at E58 from 40 animals were using UCSC conservation tracks. “.” indicate the identical sequences while “-” or “=” indicate gap in alignment.

**Supplement Figure 7.**
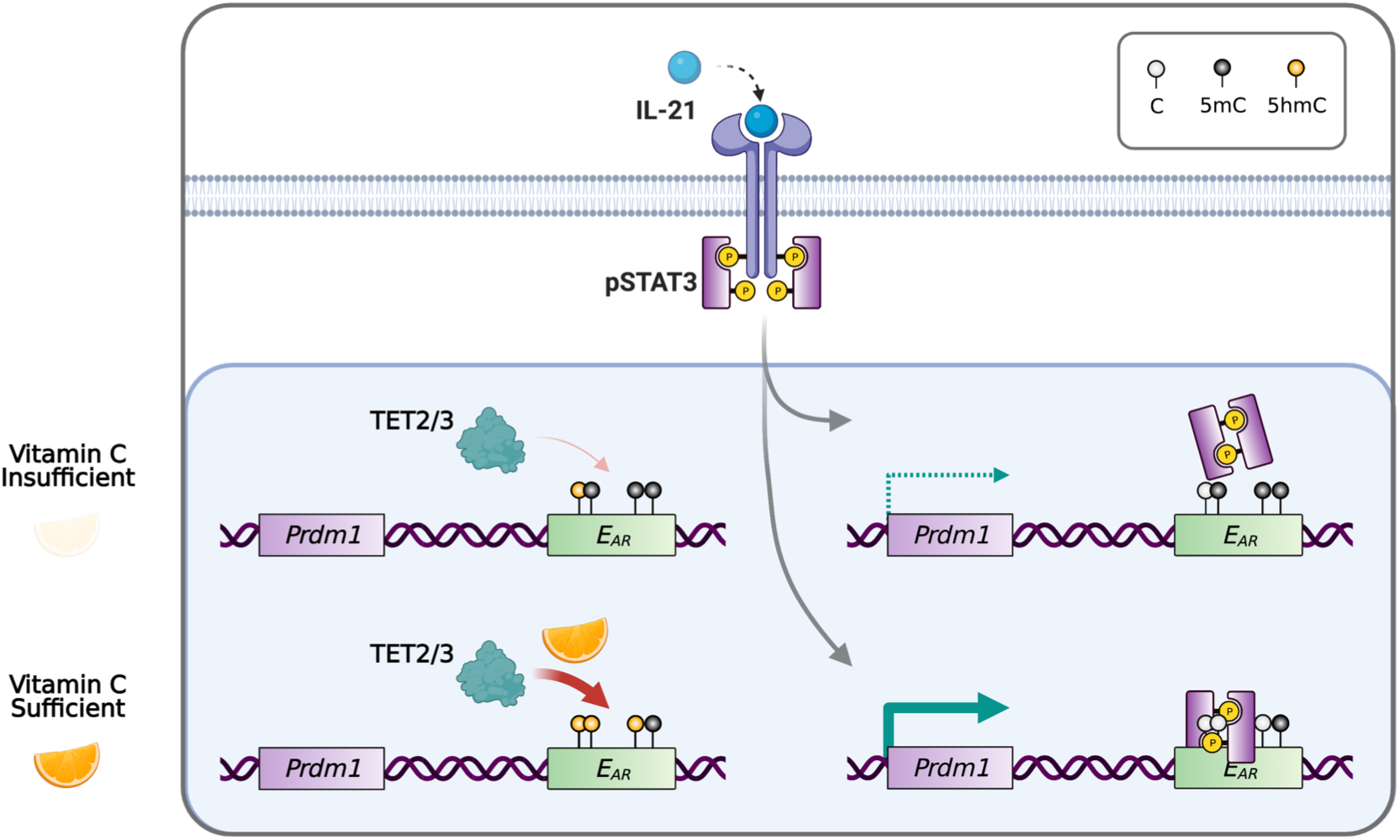
Working model for VC-facilitated plasma cell differentiation. During plasma cell differentiation, TET2 and/or TET3 is recruited to the enhancers of *Prdm1*. In the presence of sufficient VC, TET proteins have a higher enzymatic activity and efficiently oxidize 5mC into 5hmC at certain *cis* elements, which we termed “ascorbate responsive element” or E_AR_. The oxidation of 5mC at some E_AR_ may increase the association of TFs, such as STAT3 to E58. However, when VC is limited, the decreased TET activity resulted in an inefficient oxidation of 5mC, which may preclude the binding of TF and/or the maintenance the association between DNA and nucleosome (not depicted). Therefore, the activity of epigenetic enzyme may reflect the availability of micronutrients or metabolites and may influence gene expression and cell fate decisions.

## Supplementary Tables

**Table S1**. DEGs between naïve and day 4 mock B cells.

**Table S2**. DEGs between naïve and day 4 VC B cells.

**Table S3**. DEGs between day 4 mock and day 4 VC B cells.

**Table S4**. Components of B27 serum free supplement.

**Table S5**. Antibody and FACS reagent list.

